# Neural correlates of combinatorial reasoning in prefrontal cortex

**DOI:** 10.64898/2026.03.05.709889

**Authors:** Tao Hong, William R. Stauffer

**Affiliations:** Department of Neurobiology, University of Pittsburgh, Pittsburgh, PA 15213; Program in Neural Computation, Carnegie Mellon University, Pittsburgh, PA 15213; Center for the Neural Basis of Cognition, Pittsburgh, PA 15213

## Abstract

Complex economic decisions are often combinatorial: they require individuals to select from many alternatives under strict constraints on time, resources, and energy. Combinatorial reasoning is the cognitive process that enables decision makers to construct and evaluate multiple potential solutions in the face of these challenges, but its underlying neural mechanisms are not known. Using a combinatorial optimization paradigm, we show that rhesus macaques adopt distinct reasoning strategies and track the combinatorial upper bound—the best achievable outcome given the options examined so far. We recorded single units from the dorsolateral prefrontal cortex (DLPFC) - a structure central to higher cognition and complex behaviors - while the animals deliberated. Single DLPFC neurons encoded and updated the upper bound in real time, and the neuronal activity scaled with computational requirements. These findings uncover a neural correlate for combinatorial reasoning and reveal how the brain supports complex economic decisions.

## Main Text

Computational complexity quantifies the resources required to solve problems and provides formal measures of computational difficulty and algorithmic efficiency (*1*). Many everyday mental tasks — such as planning, scheduling, budgeting, inferring, and deciding — share a key feature with formally complex computational problems: they are *combinatorial*. Decision-makers engage in combinatorial reasoning when evaluating multiple options under constraints—typically those imposed by limited time, resources, or money. As the number of options decision-makers must consider multiplies, computational complexity increases. Thus, computational complexity is not merely an abstract property of formal systems, but a challenge deeply embedded in the tasks decision-makers face (*2*). And yet, the neural mechanisms for combinatorial reasoning remain unknown.

The dorsolateral prefrontal cortex (DLPFC) is widely acknowledged to be crucial for working memory and higher cognitive functions (*3*–*8*). Single unit recordings in nonhuman primates (NHP) have shown that the DLPFC is engaged during abstract categorization (*9*), probabilistic reasoning (*10*), numerical judgements (*11*–*13*), and economic decision making (*14*–*16*). In fact, it is often described as essential for complex, goal-directed behavior (*17*). Despite this, we do not know the neural mechanisms in the DLPFC that mediate combinatorial reasoning to define those goals, nor do we understand the effect and limits imposed by computational complexity - the key variable that determines the difficulty of combinatorial reasoning problems. Therefore, we recorded single neurons in DLPFC as rhesus monkeys solved problems in the *knapsack task*—a nonhuman primate (NHP) combinatorial optimization paradigm (*18*).

### Reasoning Complexity is Reflected in Visual Search

We trained two rhesus monkeys on the values of 11 items (Fig. S1). For knapsack trials, we cued trial start and item location with gray boxes, and the instance appeared after a short but random interval. Each trial lasted 5 seconds, the monkeys used a touchscreen monitor to select subsets of items, and if the sum of their selected subset was <= 0.8ml, they were rewarded with the subset sum (Fig. 1A). We classified the approach the animal used, based on the solution. As in our previous work (*18*), we classified the animals’ solutions as low- or high-complexity by matching the solution to the most likely algorithm type that generated that solution (Methods). This revealed that the animals used a mixture of low-complexity reasoning based on item value, and high-complexity reasoning based on combination values (Fig 1B, top). As previously, deliberation times corroborated our algorithmic classification scheme: that animals deliberated for longer durations when they appeared to use high-complexity algorithms (Fig. S2).

**Fig. 1.**
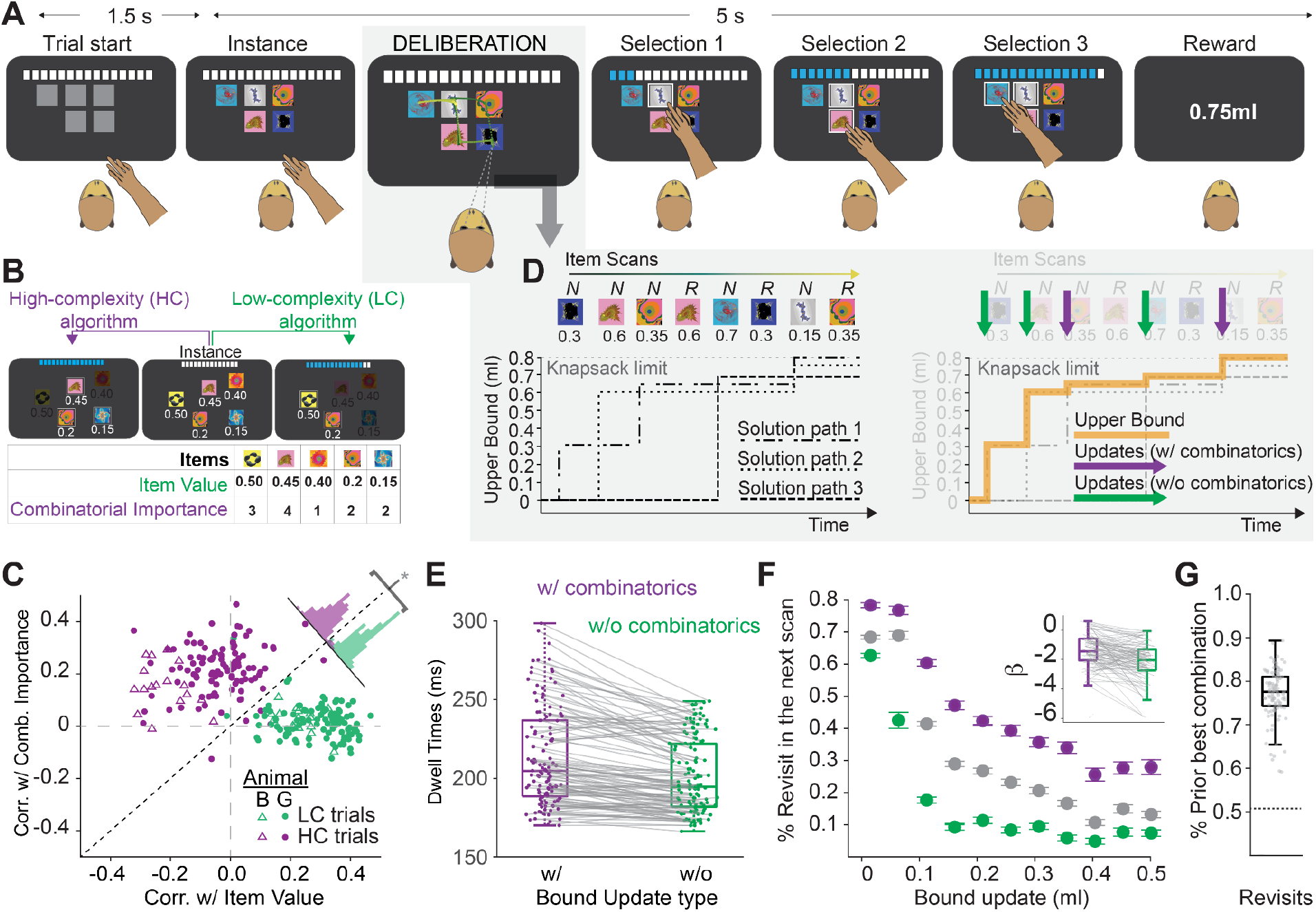
Eye movements during the Knapsack Task reflect reasoning processes. **(A)** Knapsack Task schematic. The deliberation phase is defined to be the time period between instance onset and first selection, where there is no visual feedback on the computer screen. An example eye movement trace was overlaid on top. **(B)** Low and high-complexity trials were categorized by behavioral solutions and eye search criteria (Methods). The table shows the item value and combinatorial importance of individual items in the example instance (Methods). **(C)** Scatter plots of Kendall’s rank correlation between dwell time and item value (horizontal axis) versus Kendall’s rank correlation between dwell time and combinatorial importance (vertical axis). Data from monkey G and B were shown in filled dots and triangles respectively. Correlations for low and high-complexity trials were shown in green and purple respectively. During low-complexity reasoning, eye movements were positively correlated with item value (Kendall’s rank correlation = 0.29 and 0.24, p < 3 * 10^−18^ and 3 * 10^−4^, for animal G and B, respectively, Signed-rank test) but showed minimal correlation with combinatorial importance (Kendall’s rank correlation = 0.02 and 0.03 for animal G and B, respectively, Signed-rank test). In contrast, during high-complexity reasoning, eye movements correlated positively with combinatorial importance (Kendall’s rank correlation = 0.22 and 0.16, p < 5 * 10^−18^ and 4 * 10^−4^, for animal G and B, respectively, Signed-rank test) but not with item value (Kendall’s rank correlation = −0.03 and −0.22 for animal G and B, respectively). Unity line corresponds to the dotted diagonal line. **(D)** Schematic of the evolution of the potential solution paths during the deliberation phase (left). Each path corresponds to a unique solution. The scanned items and their values were shown on top. The newly scanned items and revisits were labeled ‘*N*’ and ‘*R*’ respectively. The value of the upper bound was traced by the orange line (right). The upper bound is the best possible outcome the animal can achieve given the scan history. Purple and green arrows point to the upper bound updates that required combinatorial reasoning the updates that didn’t require combinatorial reasoning respectively. **(E)** Paired boxplots showing the dwell times for upper bound updates. Dwell times for upper bound updates that required and didn’t require combinatorics were shown in purple and green respectively (p < 10^−11^ and 0.004 for animal G and B, respectively, Signed-rank test). **(F)** Scatter plot of probability of revisiting previously scanned items in the next scan as a function of the upper bound updates (grey, β = −1.865 and −1.985 for animal G and B respectively, Logistic regression, p < 3 * 10^−18^ and 3 * 10^−4^ for animal G and B respectively, Signed-rank test). Combinatorics-required or not-required updates were shown in purple and green respectively. Error bars represent standard errors of means across occurrences of new item scans that resulted in the updates of the same magnitude. Inset: paired boxplots showing the regression coefficients of upper bound update in a logistic regression model that models the probability of revisiting previously scanned items in the next item scan (p < 5 * 10^−8^ and 4 * 10^−4^, for animal G and B respectively, Signed-rank test, Methods). Each pair of dots represent the estimated coefficients of a single session. **(G)** Boxplots showing the probability of revisiting items that belonged to the prior best combinations among all revisits (p < 10^−17^ and 10^−3^, for animal G and B respectively, Signed-rank test, Methods).

To quantify visual search behavior, we measured dwell times—the total duration the eyes remained fixated within item-specific tolerance windows—for each of the five available items during the deliberation period (Fig. 1A, shaded epoch). We performed trial-by-trial correlation analysis between eye scan path and two independent metrics: (1) item values, the learned rewards associated with the items, and (2) combinatorial importance, a measure of how often an item appeared in ‘good’ combinations (Methods). Dwell times were positively correlated with item value during low-complexity reasoning trials (Fig. 1C, green), and combinatorial importance during high-complexity reasoning trials (Fig. 1C, purple). These distinct correlations indicate that the animals employed search strategies that reflected different reasoning complexities (Fig. 1C, inset histograms). These results complement our previous findings (*18*), and provide the strongest evidence to date that the animals use distinct psychophysical processes, analogous to computational algorithms, to search for satisfactory solutions.

### Upper Bound Guides Eye Search Process during the Knapsack Task

The ‘upper bound’ is a variable computed by combinatorial algorithms and used to measure the progress of optimization and to serve as a benchmark for evaluating different combinations (*19*–*22*). Here, we sought to examine whether the concept of the combinatorial upper bound was relevant to the animals’ behavior. As the animals made saccades, different combinations of items could be used to form combinations and potential solutions (Fig. 1D, left, hashed lines). The upper bound is defined as the maximum value, over all potential solutions (Fig. 1D bottom, Orange line). Note that, upper bound increases could be achieved by updates that required combinatorics and those that did not (Fig. 1D, bottom, purple vs green arrows). We found that the animals dwelled longer on items that generated bound updates via combinatorics, compared to items that generated bound updates that did not require combinatorics (Fig. 1E). This result is consistent with combinatorial bound updates requiring more computations. Furthermore, we found that the probabilities of the animals scanning a new item or revisiting a prior scanned item varied according to the type and magnitude of bound updates. When the bound updates were small, the animals were more likely to revisit the previously scanned items (Fig. 1F, grey). This effect was stronger when the updates required combinatorics to achieve, vs. when they did not require combinatorics (Fig. 1F, 1F inset, purple vs green). This result suggests that the animals might revisit items for comparing the values of the combinations that are close in value. To test the idea that revisitations reflected comparisons, we tested whether the animals were more likely to revisit items that belonged to the prior best combinations. We found that the animals revisited items that belonged to the prior best combinations around 77% of the time, significantly above chance (Fig. 1G). These results demonstrate that the combinatorial variable ‘upper bound’ is reflected in the behavior, and that eye movements reveal discrete steps of the deliberation process as well as their complexity.

### Upper Bound Representation in the Dorsolateral Prefrontal Cortex

We recorded 202 neurons from the DLPFC of two rhesus macaques performing the knapsack task (Fig. 2A, n = 101 in animals B and G, each). We aligned single neuron impulses to saccade onsets and analyzed the neural response to bound updates during the deliberation period (Methods). Saccades to items that improved the upper bound evoked responses that reflected closely the magnitude of the upper bound changes (Fig. 2A-D, left). We discovered that 37 single neurons - 23 and 14 in animal G and B, respectively - modulated their impulse rates monotonically as a function of the bound updates (Fig. 2H-I). The vast majority (28/37) of these ‘bound update neurons’ showed no significant responses following saccades that did not increase the upper bound (Fig. 2A-D, right) and only a small subset (3/37) generated responses modulated by saccade direction (Fig. S3). Since there was no visual feedback during the deliberation period, the bound-update neural responses also did not reflect physical changes on the screen. These results show that single DLPFC neurons encode discrete computations of the combinatorial upper bound, and demonstrate a potential neural substrate of combinatorial reasoning in DLPFC.

**Fig. 2.**
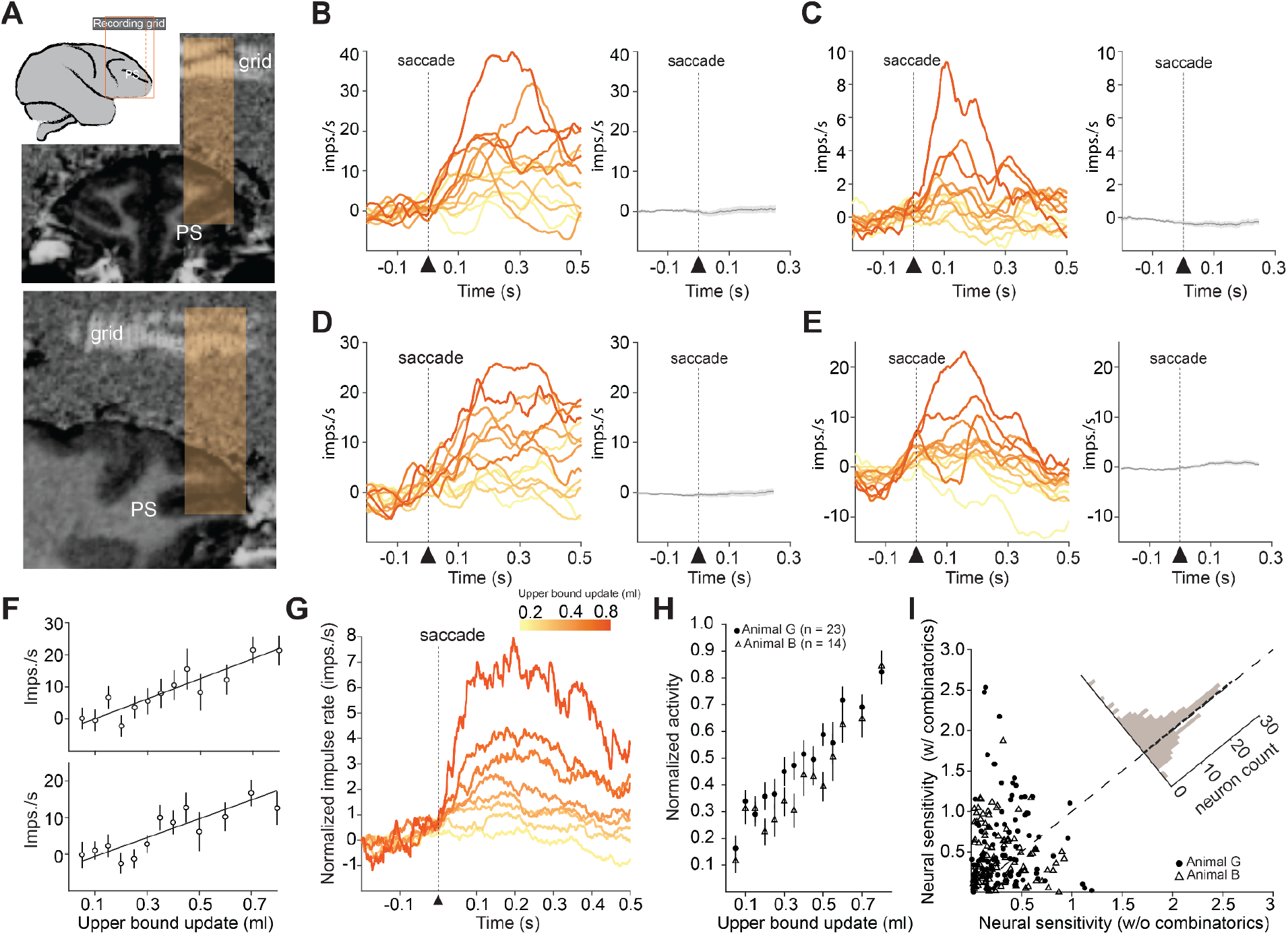
Single neurons in DLPFC encode upper bound updates. **(A)** (*inset*) Schematic of macaque brain: the dashed line indicates the approximate coronal plane in top MRI, whereas the orange square indicates the region shown in the bottom, sagittal MRI. (*top*) Image of coronal MRI scan. The recording grid was filled with petroleum and visualized as white lines in the MRI scan. The shaded rectangles indicate recording angles and locations. (*bottom*) as in top, for sagittal view of MRI scan. **(B-E)** Peri-event histograms (PETH) of four example bound-update neurons in DLPFC. The left PETHs were aligned to the onsets of saccades that led to upper bound increases. Neural activity traces for upper bound changes are colored by the corresponding magnitude. Scale located above panel G. The right PETHs were aligned to onsets of saccades that led to no changes in the upper bound. Shaded areas are standard errors of means. A and C depict two neurons recorded from animal G. B and D depict two neurons recorded from animal B. **(F)** Scatter plots of impulse rate vs mound update magnitude for two example bound-update neurons in DLPFC (ρ= 0.901 and 0.846, p < 10^−4^ and 0.0009, respectively, Spearman’s correlation). Error bar represents standard errors of mean across saccades that led to the same improvements of upper bound. **(G)** Population-averaged PETH of all bound-update neurons. Activity traces are binned into eight bins and are colored based on the update magnitude. **(H)** Scatter plots of bound-update neural population averages for animal G (filled dots) and B (triangles). Error bars represent standard errors of mean across neurons (ρ= 0.982 and 0.903, p < 10^−8^ and 10^−6^, for animal G and B, respectively, Spearman’s correlation). **(I)** Scatter plot of neural sensitivities to upper bound changes from every neuron. Vertical and horizontal axes show the neural response slopes when combinatorics was or was not required, respectively, to improve the upper bound. Neurons recorded from animal G and B are shown in filled dots and triangles respectively (p < 10^−4^ and 10^−3^ for animal G and B, respectively, Signed-rank test). Inset: the histogram shows the density of the neurons across both animals relative to the unity line (dotted line).

For each neuron, we separated responses according to the complexity of the associated upper bound updates - those that required combinatorics and those that did not. The majority of neurons exhibited greater sensitivity to changes in the upper bound when high-complexity combinatorics were required (Fig. 2J). This result demonstrates that the complexity of ongoing computations modulated the activity of DLPFC neurons.

In light of the evidence of phasic responses reflecting bound updates, we examined ramping activity in single neurons to look for coding of the upper bound itself. We used Spearman’s correlation to screen for neurons that reflected the upper bound (Fig. 3A-B, insets). After False Discovery Rate (FDR) correction, this analysis revealed that 32 neurons were significantly modulated by the upper bound (p < 0.05, Benjamini-Hochberg Procedure). Since the upper bound growth rate is dependent on the items and the animals’ visual search, the upper bound grew faster on some trials than others. Therefore, we aligned neural activity to the first scanned item and separated trials according to high and low rates of upper bound growth (Methods). The PSTHs of the significant neurons demonstrated monotonically increasing impulse rates that reflected the different rates of growth in the upper bound (Fig. 3A-C). To quantify this relationship, we modeled the evolution of neural impulse rates during the deliberation phase as a linear function and used the estimated trial-wise slopes to characterize the buildup of the neural activities since the first item scans (Methods). The trial-wise upper bound growth rates correlated positively with the neural buildup rates: upper bound neurons show steeper ramping slopes during trials with faster upper bound growth (Fig. 3D). To control for potentially correlated variables, such as specific movement preparations or general preparatory signals, we regressed the neural impulse rates of these 32 neurons against these signals, and then computed Spearman’s correlations between the upper bound and the residuals in the impulse rates (Methods). This analysis demonstrated that 22 of 32 neurons reflected upper bound even after controlling for correlated variables (p < 0.05, Spearman’ s correlation). These results demonstrate that a subset of ramping DLPFC neurons code for the combinatorial upper bound, and that this activity was not mediated by other, more generic preparatory processes.

**Fig. 3.**
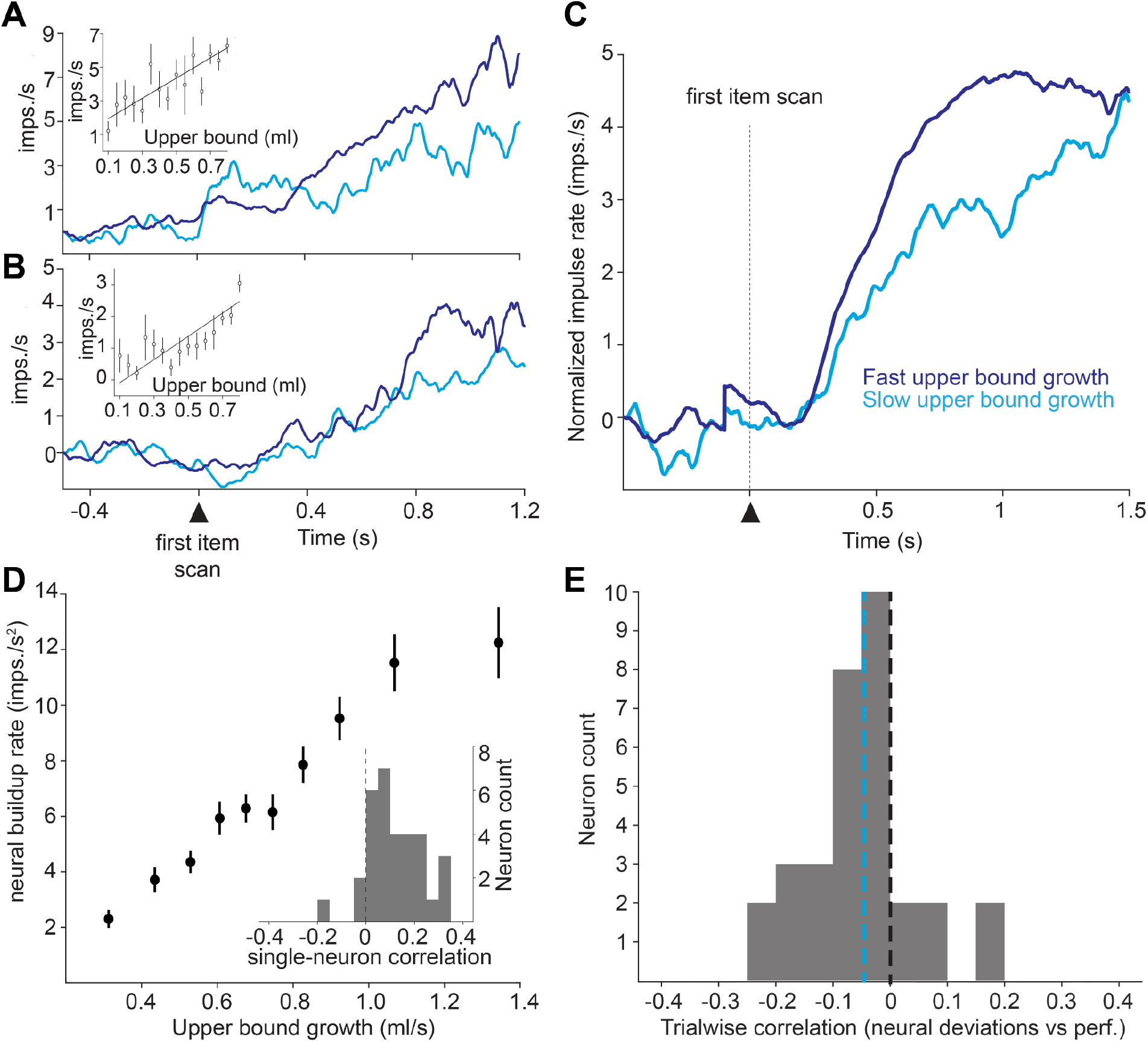
Single neurons in DLPFC track the upper bound. **(A-B)** Peri-event histograms (PETH) of two example upper bound neurons in DLPFC. PETHs are aligned to the start of the first item scan in the trial. Trials are grouped based on upper bound growth rate (<= 0.6 ml/s or > 0.6ml/s, Main text). Average activity traces for fast upper and slow bound growth trials are in dark and light blue respectively. Insets: scatter plots showing the average neural activities of the corresponding neuron as a function of upper bound (ρ= 0.839 and 0.782, p = 0.0001 and 0.0009, A and B, respectively). Error bars are standard errors of means across the same upper bound values. **(C)** Population-averaged PETH of all upper-bound neurons recorded from animal G. **(D)** Scatter plot showing the average neural buildup rate as a function of upper bound growth. Trials were binned into ten equal sized bins based on upper bound growth rate. The median upper bound growth rate of each bin was used. Error bars represent standard errors of means across trials within the same bin. Inset: Histogram showing the Spearman’s correlation between neural building rate and upper bound growth for individual upper bound neurons (p < 10^−4^, Signed-rank test). **(E)** Histogram showing the trial-by-trial Spearman’s correlation between upper-bound related impulse rate deviations and the rewards the animals received at the end of the trial (p = 0.0045, Signed-rank test, Methods).The blue dotted line corresponds to population-average correlation.

If the DLPFC actively participates in combinatorial reasoning by computing and representing the upper bound, upper-bound-related neural variability should be related to performance of combinatorial reasoning. In other words, larger deviations from average upper-bound impulse rates should correlate with lower performance on a trial-by-trial basis. For each Upper Bound Neuron, we computed the trialwise impulse rate deviations and correlated with the rewards the animal received (Methods). The Upper Bound neural population showed negative correlations between neural variability and rewards (Fig. 3E). This trial by trial correlation between upper bound coding and combinatorial reasoning performance suggests an active role for DLPFC in combinatorial reasoning.

### Bound Update and Upper Bound neuron populations were largely distinct

In computational models, the upper bound changes according to bound updates, and thus we wondered if Upper Bound Neurons were a subset of Bound Update Neurons. Surprisingly, we found that the fraction of neurons classified as both Upper Bound and Bound Update Neurons were small and did not differ from chance level (Fig. 4A). To better understand how a neuron could be an Upper Bound Neurons and not a Bound Update Neuron, we aligned the Upper Bound Neurons to the onsets of item saccades that led to bound updates, and plotted population PSTHs. This analysis showed that Upper Bound Neurons were slowly increasing around upper bound changes (Fig. 4B, blue), but did not respond with the same phasic responses as Bound Update Neurons (Fig. 4B, orange). Intriguing, the average dwell time on single items corresponded to the intersection of activities between Bound Update and Upper Bound neurons. This result suggests that Bound Update and Upper Bound neurons could communicate to facilitate combinatorial reasoning. Altogether, these results demonstrate two distinct functional populations of DLPFC neurons that contribute to combinatorial reasoning and suggest a neural circuit implementation for these complex deliberations.

**Fig. 4.**
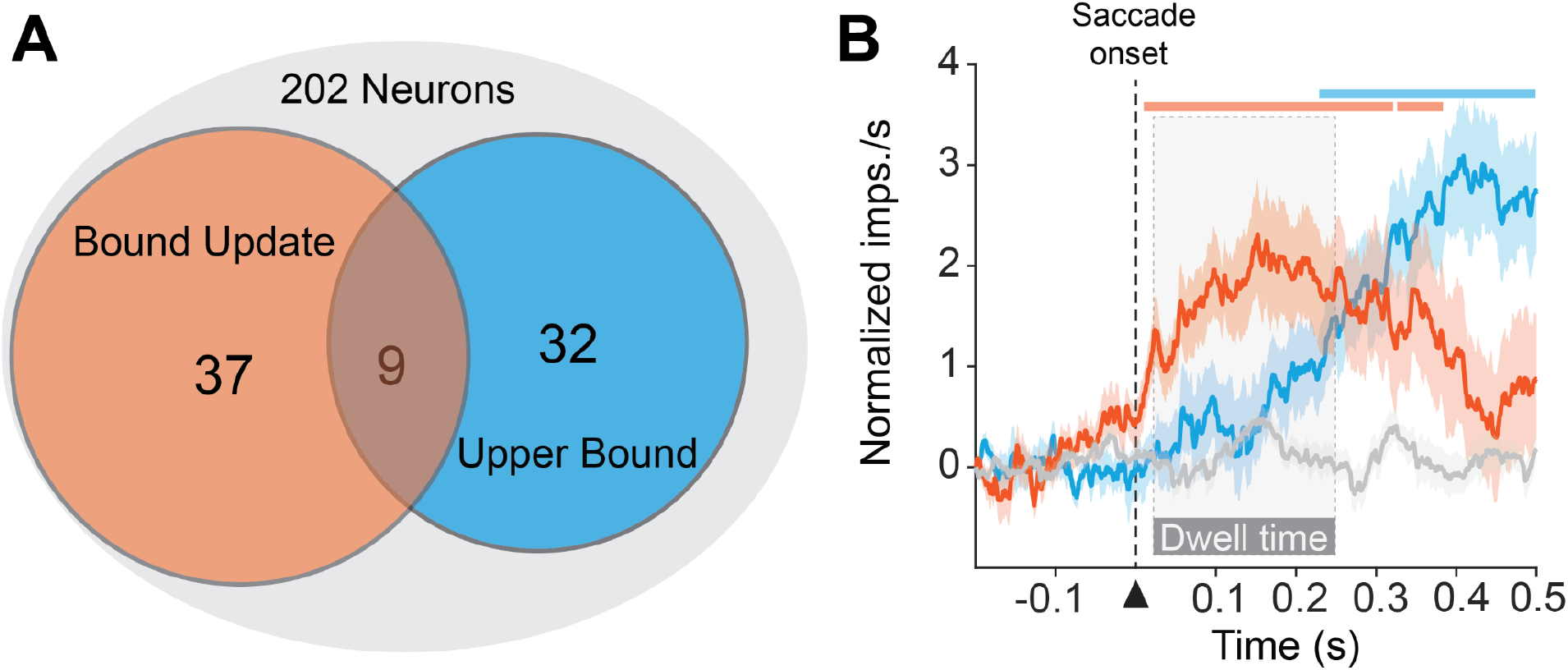
A potential neural circuit for combinatorial reasoning. **(A)** Venn diagram showing the total number of recorded DLPFC neurons, upper bound neurons, and bound update neurons in grey, blue and orange, respectively (not the scale). The overlap between Upper Bound and Bound Update Neurons did not differ from chance level (p = 0.136, chi-squared test). **(B)** Population PETHs showing the neural dynamics of the bound update subpopulation (orange), upper bound subpopulation (blue) and the rest of the recorded neuron (grey). Shaded areas correspond to s.e.m across neurons. Orange and blue bars represent the time period when the activity of the bound-update and upper-bound population significantly differed from 0, respectively (p<0.05, Signed-rank test, Benjamini-Hochberg Procedure). The mean dwell time window relative to saccade onset was marked by a partially transparent rectangle. Only segments longer than 20ms were shown.

## Conclusions and Discussions

Here we adapted the knapsack task to enable behavioral neurophysiology and eye tracking. At the level of single trials, the animals’ visual search patterns reflected the complexity of their thought process (Fig. 1C). At the level of single saccades occurring during deliberation, the duration of fixation on individual items reflected the computational complexity of calculations needed to update to the combinatorial upper bound - the best possible solution given all items scanned (Fig. 1D,E). Moreover, when the animals made incremental improvements to upper bound, and even more when those updates that required combinatorics to achieve, they revisited items from the prior best solution, suggesting that they were comparing combinations. Altogether, these psychophysical data demonstrate algorithm-like reasoning behavior in the face of computational complexity. We recorded single unit activity from the DLPFC (Fig. 2A). Single neurons were phasically active following saccades that led to upper bound improvements, and the magnitudes of those responses reflected the magnitude of the change in upper bound (Fig. 2B-I). Single neurons were more active when these bound updates required combinatorics (Fig. 2J), consistent with the concept that higher complexity computations require more operations. We also found neurons that coded the level of the upper bound, and the fidelity of their upper bound coding predicted the rewards the animals received on a trial-by-trial basis (Fig. 3). These two functional types were largely distinct, rather than overlapping populations (Fig. 4), and this fact suggests a recurrent neural circuit for finding good solutions to complex problems. In this circuit, the upper bound neurons, with their longer memory trace, would track the current best outcome, and serve as a baseline for bound update neurons. In turn, bound updates neurons, when they are phasically active, would drive the activity of upper bound neurons towards a higher upper bound. Together, these two neuron types provide signals that are crucial components of any information processing system that needs to achieve combinatorial reasoning. Thus, our results demonstrate neural correlates of combinatorial reasoning. In the context of value optimization, these results indicate that the DLPFC contains the neural machinery to aid decision makers in determining what they want in the face of computational complexity.

Decision makers routinely encounter situations with high computational complexity, including budgeting, managing households, and socializing. In contexts like these, combinatorial reasoning is used to define priorities that exist in large solution spaces and that satisfy constraints such as time, monetary resources, and energy levels. Notably, patients with prefrontal dysfunction are impaired on tasks with high computational complexity or that require combinatorial reasoning. For example, individuals with schizophrenia exhibit marked deficits in tasks requiring combinatorial reasoning, and these deficits are stronger predictors of long-term prognosis than the severity of psychotic symptoms (*23, 24*). Household wealth—a proxy for budgeting ability—starts to decline well before a dementia diagnosis (*25*). People in the preclinical stages of Alzheimer’s disease have higher rates of delinquency on mortgage payments than age- and sex-matched controls (*26*). Many of these same deficiencies appear in the prodromal phase of Parkinson’s disease and are predictive of Parkinson’s disease dementia (PDD) (*27*). Therefore, understanding the neural basis of combinatorial reasoning is crucial to understanding some of the most debilitating mental health disorders, and our results are a crucial step in that understanding.

We used computational methods for combinatorial reasoning as our basis for examining the neural basis of combinatorial reasoning. A high-level concept that is common to many of these algorithms is the combinatorial upper bound (*19*–*22*). We operationalized the upper bound by linking it to visual search, and defined as the most valuable outcome possible, computed over all the items the animals had scanned to that point. This framework represents a significant divergence from previous studies of value-based decision-making in monkeys. In many studies, value is dictated by presented cues or learned through foraging (*28, 29*). In contrast, the upper bound cannot be learned via reinforcement. In the knapsack task, the values of the individual items remained unchanged, but the trajectory of the upper bound varied across trials. The same item can elicit upper bound improvements of different magnitude depending on the items the animals previously scanned. Thus the upper bound must be computed in real time. In this regard, the bound update neuronal responses shown here have some parallels with dopamine reward prediction error responses. The magnitudes of prediction errors and dopamine prediction error responses are determined by the differences between received and predicted rewards and calculated in real time (*30, 31*). However, reward prediction errors are learning signals (*32, 33*), and likely have small influence over ongoing decisions. The upper bound, in contrast to reward prediction error, has no known role in learning but, as we show here, is constructed and actively used to determine the most desirable outcomes. Moreover, the computational requirements of upper bound updates influence the animals’ behavior even after controlling for their magnitudes, which are consistent with the idea that computationally demanding economic decisions should be understood through the lens of computational complexity theory (*34*).

Reasoning is crucial to effectively navigating the world, and there are many forms of reasoning. We previously showed that the combinatorial reasoning in the knapsack task can last seconds (*18*), and the upper bound neurons track the upper bound across multiple bound updates during that time period. In this aspect, upper bound coding resembles neural activities during probabilistic reasoning where single neurons in DLPFC, OFC and LIP integrate probabilistic evidence for or against a decision over time (6,33,34). However, probabilistic reasoning is fundamentally distinct from combinatorial reasoning. Probabilistic reasoning allows decision makers to cope with unpredictability and infer hidden structure from stochasticity, whereas efficient combinatorial reasoning facilitates searching a deterministic—but often exponentially large—solution space to identify feasible or optimal solutions under computational constraints. In other words, the factor that makes probabilistic reasoning challenging is ambiguity, whereas the factor that makes combinatorial reasoning challenging is computational complexity. This fact is reflected in the tasks used to study the neural correlates. In probabilistic reasoning, hard trials are trials with less probability information, whereas in the knapsack task, hard trials are trials that require more computational resources to solve. As a result, upper-bound related neural activities reflect those computations: the constructions, evaluations and comparisons of potential solutions. Furthermore, the fact that the coding of upper bound updates is well aligned to item saccades suggests that upper bound coding is a neural correlate of a discrete, algorithmic reasoning process. Despite these differences, together these studies highlight a network in the brain responsible for reasoning and intelligence.

During preliminary testing for the neurophysiological recordings, we discovered that instance repetition quickly eroded the deliberation times. Because we were recording with one or a few electrodes at a time, we chose ten instances from the available 462, and showed them repeatedly to ensure we had enough repetitions to construct neural statistics. However, the resulting data showed that within a session, the deliberation times steadily decreased with every instance repetition. It appeared that the animals quickly learned the ‘right’ answer to a given problem, and from that point forward, they no longer had to think, just act, in order to get good rewards. Therefore we discarded that approach, and reverted to the approach we had used for our prior behavioral study: pseudorandomly selecting instances from all available instances. The animals did 250-300 trials per day, and this mostly prevented the animals from experiencing the same instance twice in one day. This approach limited us from examining neural correlates of solutions to particular instances. We instead focused on the upper bound and bound updates, as they are concepts that span all instance types. In future studies, we will use pixel-style recording arrays to gather many neurons simultaneously and examine the evolution of single reasoning processes within neural ensembles.

Despite these limitations, we were able to show that DLPFC neurons code for a particular combinatorial value signal, the upper bound. Our data indicate that the DLPFC contains a circuit of distinct functional neuron types, one that codes for the level of the upper bound and another that codes changes in the upper bound. We showed that the fidelity of upper bound coding was directly related to the animals’ performance. The DLPFC has been implicated in the neurobiology of sophisticated cognition ever since the seminal discoveries that DLPFC neurons are active during the delay period of working memory tasks (*6*–*8, 35*). It is also known that patients with DLPFC dysfunctions struggle with tasks that have high computational complexity (*23*–*25, 27*). Our findings extend the function of DLPFC neurons to combinatorial reasoning, and directly link their functions with DLPFC dysfunctions.

## Acknowledgments

We thank Katalin M. Gothard for discussion and critical reading of the manuscript. We thank Jackie Breter for animal care.

## Funding

National Institutes of Health grant DP2MH113095 (WRS)

National Institutes of Health grant R01MH128669 (WRS)

## Author contributions

Conceptualization: TH, WRS

Methodology: TH, WRS

Investigation: TH, WRS

Visualization: TH, WRS

Funding acquisition: WRS

Project administration: WRS

Supervision: WRS

Writing – original draft: TH, WRS

Writing – review & editing: TH, WRS

## Competing interests

Authors declare that they have no competing interests.

## Data and materials availability

All data are available in the main text or the supplementary materials

## Methods

### Animals, Surgery and Setup

All animal procedures were approved by the Institutional Animal Care and Use Committee of the University of Pittsburgh. We used two male rhesus macaque monkeys (*Macaca mulatta*) for these studies (one 12 years of age, one 8 years of age, 12.4kg and 9.5 kg). A titanium head holder (Gray Matter Research) and a recording chamber (Crist Instruments, custom made) were aseptically implanted in one animal under general anesthesia before the experiment. A custom plastic head holder and recording chamber (Rogue Research) were aseptically implanted in another animal under general anesthesia before the experiment.

The recording chamber for vertical electrode entry was centered 35 mm anterior to the interaural line for one animal and 38mm for the other. During experiments, animals sat in a primate chair (Crist Instruments). During behavioral training, testing and neuronal recording, eye position was monitored using infrared eye tracking (Eyelink Plus 1000). Custom-made software (Matlab, Mathworks Inc.) running on a Microsoft Windows 7 computer controlled the behavioral tasks. We used custom-made software and Spike 2 (CED 1401) to collect behavioral and neural data.

### Behavioral Task

Each stimulus set contained 12 fractal images that predicted rewards between 0.1 and 0.8 ml. We trained the animals on the predictive value of each of the 12 fractal images. On each training trial, the screen presented a grey square that indicated the position of the item would be shown, and the virtual knapsack. After a 1.5s delay, one item was pseudorandomly selected from the set and replaced the grey square. When the animal touched the screen where the item was presented, the virtual knapsack was filled with the ‘volume’ associated with the item, and the animals were rewarded with that same reward volume. Training for each stimulus set lasted one month. We used the animals’ response times to measure learning progress (Fig. S1).

Each knapsack trial began with five grey squares that indicated the positions of the items. Their locations were pseudorandomly selected from the six potential locations. After a 1.5s delay, an ‘instance’ - a combination of five items that appeared, simultaneously, on the screen. The animals used the touchscreen to select items one-by-one. Every time an item was selected two things happened, (1) that item was highlighted and (2) the virtual knapsack at the top of the screen filled by an amount equivalent to the item’s predicted reward. The knapsack limit was set at 0.8 ml. This limit was chosen based on prior studies that demonstrated the reward utility functions were relatively linear around 0.8 ml. Optimal performance on every trial was defined as the largest possible sum of rewards less than or equal to the limit. If the monkey selected a combination of items whose reward magnitude sum was less than optimal, they were rewarded by that lesser amount. If they selected a combination that was greater than the limit, then the trial ended, no reward was delivered.

### Analysis of Behavioral Data

#### Matching algorithms to behavior solutions

This method was described in detail in our prior publication (*18*). For each trial, we first calculated the graph distance between the solutions generated by three candidate algorithms - the greedy (*36*), Sahni-k (*37*), and Johnson-t algorithms (*38*) - and the animals’ solutions. We constructed an undirected graph, where the nodes of the graph represented all potential combinations. Two nodes are connected if one node can be reached from another by adding or removing a single item and the corresponding edge weight is the value of that item (*39*). We defined the distance between any two combinations to be the length of the shortest path between the two representative nodes. The distance between any two nodes *n*_1_ and *n*_2_ can be computed as follows:

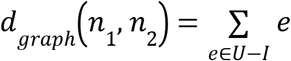

where *union*(*U*) − *intersection*(*I*) refers to set subtraction between set *U* and *I* and *e* refers to an element in the set subtraction. Note that for graph distance, the order of the items is not considered. Among the algorithmic solutions that have the smallest graph distances to the behavioral solution, we then ranked the candidates according to L1 distance. Each solution was defined as an ordered tuple and padded with zeros. For example, if the monkey selected 0.4 ml, then 0.2 ml, then 0.1 ml and then stopped. The solution was defined as (0.4, 0.2, 0.1, 0, 0). Thus, the distance between two solutions *p* and *q* can be characterized by the L1 distance.

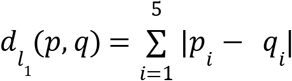

Since the high-complexity algorithms don’t specify selection order for the items considered during combinatorial search, we ordered the items selected during combinatorial search to yield the smallest L1 distance to the behavioral solution.

To minimize the possibility that all algorithms match the behavioral response poorly, we constructed a null distribution for each trial by calculating the L1 distance between all possible solutions (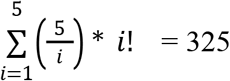 in total) and the behavioral response. The best-matching algorithm has to have a smaller L1 distance to the behavioral solution than the lower 5% threshold of the null distribution. Trials where this criterion was not met were labelled as ‘unclassified’. Finally, we denoted all solutions classified as greedy as ‘low complexity’ reasoning, and combined all solutions classified as Sahni-k or Johnson-t and denoted them as ‘high-complexity’ reasoning. These results were further refined using visual search behavior, described in the next section.

#### Analysis of visual search behavior

We defined a region of interests (ROI) surrounding the presented items and only gazes that fell inside of the ROI were used for analysis. An item scan was established if the gaze remained in the ROI for at least 100 ms.

##### Dwell time analysis

To compute the total dwell time of an item, we summed the durations of all corresponding item scans during the time between instance onset and the animals’ last selections. Let *CoI* _*i,S*_ denote the combinatorial importance of an item *i* within instance *S*. Let *G*_*S*_ be the set of ‘good’ solutions: solutions whose sums are greater than or equal to the animals’ median performance for instance *S*. Then, the combinatorial importance for item *i* within instance *S* can be defined as follows:

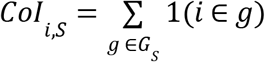

For each trial, we then computed Kendall’s tau-b correlation between dwell time and item value or combinatorial importance separately and averaged the correlation coefficients across trials within the same sessions. Trials were categorized into low- and high-complexity trials first based on a previously developed classification scheme (Hong and Stauffer, 2023). Leveraging eye tracking data, we were able to perform more stringent trial classifications as follows. It is possible that the animals generated solutions resembling those generated by high-complexity algorithms via a step-by-step process instead of combinatorial search, or vice versa. One of the key differences between combinatorial and step-by-step search algorithms in the knapsack task is whether the selected items were considered as combinations before the first selection. The eye movement data allowed us to incorporate an additional criterion to reflect this difference: the animals must scan more than one selected item before the first selection and no more than one selected item after the first selection. As a result, we ended up with four categories of trials: true ‘high-complexity’ trials, ‘high complexity’-like trials, true ‘low-complexity’ trials and ‘low-complexity’-like trials, as specified below. We only used true ‘high-complexity’ trials and true ‘low-complexity’ trials for the dwell time analysis. Using the base classification scheme generated similar results.

#### Upper Bound Definition

During the deliberation period, let the sequence of items the animal scans to be (*x*^1^, *x*^2^, *x*^3^, …,*x* ^*T*^), where *x*^*t*^ is the *t*-th item the animal scans. Let the set of unique items the animal had scanned up to item scan *t* to be *Y*^*t*^ = {*y*_1_, *y*_2_, …, *y*_*n*_}. We then computed the upper bound as the best possible reward achievable based on the items the animals had scanned so far in a trial. The upper bound at item scan *t* can be defined formally as:

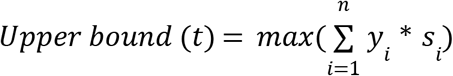

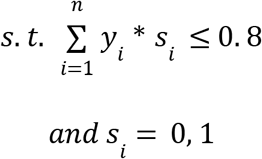

where *s*_*i*_ = 1 indicates that item *y*_*i*_ is included in the combination that achieves the current upper bound at item scan *t*. The change in upper bound at item scan *t* can be subsequently defined as:

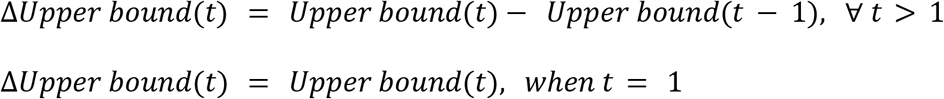

#### Upper Bound Updates and Item Revisitations

For each session, we were interested in the probability that the animal was going to scan an item that was previously visited following an item scan *t*:

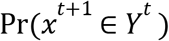

and use logistic regression to model the influence of upper bound updates on the animals’ behavior:

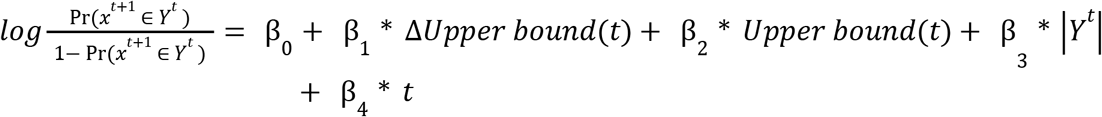

where β_1_ quantifies this influence for each session and we used a Signed-rank test to test whether session-averages are significantly different from 0. The variable |*Y*^*t*^|corresponds to the number of unique items the animals had scanned up to item scan *t*. This variable is included because by chance the animals are more likely to scan an item that was previously visited as they scan more items during deliberation. Restricting the analysis to only new item scans (i.e. modeling Pr(*x*^*t*+1^ ∈ *Y*^*t*^ | *x*^*t*^ ∉ *Y*^*t*−1^)) produced quantitatively similar results as reported in the main text (β_1_ = −2.159 and −1.91 for animal G and B respectively, Logistic regression, p < 4 * 10^−18^ and 3 * 10^−4^ for animal G and B respectively, Signed-rank test).

Let *C*^*t*^ be the current best combination - the combination that achieves *Upper bound* (*t*). For each revisitation following a newly scanned item, we also determined, for each session, the probability that it was made to an item that was part of the prior best combination:

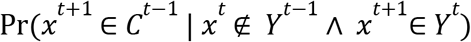

We matched the exact same sequence of item scans prior to the scan and randomly selected an item from the scan history to calculate the chance level of scanning an item that was part of prior best combinations. A Signed-rank test was performed to compare the data with chance level across sessions.

As highlighted in Fig. 1, there are two types of upper bound updates: (1) upper bound changes where combinatorics is not needed: the newly scanned item alone is greater than the previous upper bound value:

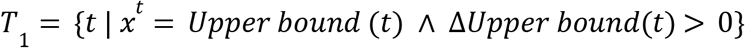

and (2) upper bound changes where combinatorics is needed: the new upper bound can only be achieved by adding the newly scanned item to the items in the prior scan history:

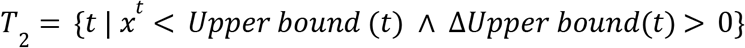

We extended the logistic regression model above to examine the different effects of these two types of upper bound updates:

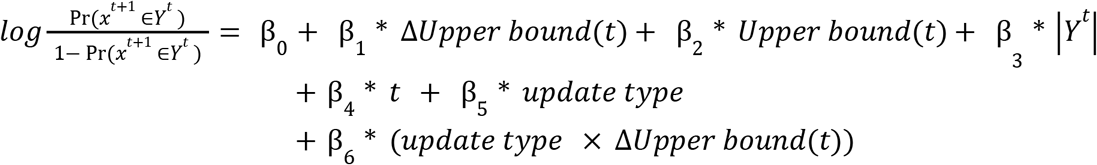

where β_6_ quantifies the interaction effect between upper bound update types (i.e. *t* ∈ *T*_1_ or *t* ∈ *T*_2_ modeling) and the magnitude of the updates, where bound updates that didn’t require combinatorics were set to be the reference level in the model. Restricting the analysis to only new item scans (i.e. Pr(*x*^*t*+1^ ∈ *Y^t^* | *x^t^* ∉ *Y*^*t*−1^)) produced quantitatively similar results as reported in the main text (β_1_ = −2.369 and −3.582, β_6_ = 0.851 and 3.508 for monkey G and B respectively, Logistic regression, p < 3 * 10^−7^ and 3 * 10^−4^ for monkey G and B respectively, Signed-rank test).

### Neural Data Acquisition

We used MRI to localize the placement of dorsolateral prefrontal cortex recording chambers. Recording grids were also visualized in MRI post-surgery using petroleum to guide placements of the electrodes. Post-surgery MRI and electrical stimulations were used to localize the dorsolateral prefrontal cortex (DLPFC) with respect to the frontal eye field (FEF). For FEF stimulations, we used electrical stimulation consisting of 100-ms trains of biphasic current pulses (200 Hz). Current amplitude was measured via the voltage drop across a 1kΩ resistor in series with the return lead of the current source. The FEF was identified by our ability to evoke saccadic eye movements with stimulation using currents around 50 - 75 μA (*40*). For animal B, DLPFC recordings were taken 35mm to 39mm anterior to the interaural line. For animal G, recordings were taken 35mm to 41mm anterior to the interaural line. Microelectrodes (FHC or AlphaOmega) or Plexon 16-channel S-probes were positioned inside a stainless-steel guide cannula and advanced by an oil-driven micromanipulator (Narishige MO-97). Action potentials from single neurons were amplified, filtered (bandpass 300 Hz to 6 kHz), and converted into digital pulses when passing an adjustable time–amplitude threshold (Bak Electronics), or sorted offline using Plexon Offline Sorter. We stored both analog and digitized data on a computer using custom-made data collection software (MATLAB) and Spike2.

### Analysis of Neural Data

#### Bound update neurons

To identify bound update neurons, we first detected saccades by thresholding eye movement speed (the speed had to be greater than equal to 20°/s for more than 10ms). For item saccades that led to changes in the upper bound, we then isolated neural response changes by subtracting the impulse rates 250ms prior to saccade onsets from the average post-saccadic impulse rates. For animal G, we reported results using average impulse rate taken in the time window after the saccade offsets and before the onsets of the saccades that led to the next upper bound change. Using a fixed time window after saccade offsets (ranging from 350ms to 500ms) generated similar results. For monkey B, we used a fixed time window of 250ms after saccade offsets. Finally, we correlated the magnitude of the upper bound changes with their corresponding, mean impulse rate changes (Spearman’s correlation, p < 0.05). We excluded upper bound updates with less than five data points. To compute population averages for bound update neurons, we normalized each neuron’s firing rates to between 0 and 1:

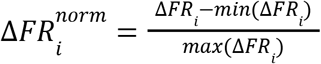

where Δ*FR*_*i*_ is the average change in impulse rate for upper bound improvement *i*. For neurons that showed negative encoding of bound updates, we flipped their normalized activities before averaging. We excluded upper bound updates with less than two neurons when computing population averages.

We also assessed whether saccade directions modulated bound-update coding by first assigning each saccade into one of eight directions (from 0 degree to 360 degrees with steps of 45 degrees) and used a two-way ANOVA to assess if the interaction term between bound update and saccade direction was significantly different from zero. For saccades that did not lead to upper bound changes, we used saccades that were sufficiently distant in time from saccades that elicited changes in the upper bound (>= two saccades). Average impulse rates −250ms to 0ms prior to saccades were compared to the average impulse rate during 0ms to 250ms after saccades with a Signed-rank test.

#### Neural sensitivity

To estimate how sensitive the average neural response changes were to changes in the upper bound, we use simple linear regression:

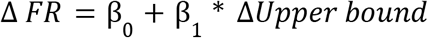

We estimate neuronal slopes, 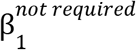and 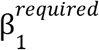, separately for two distinct types of upper bound changes: (1) upper bound changes where combinatorics is not needed: the newly scanned item alone is greater than the previous upper bound value and (2) upper bound changes where combinatorics is needed: the new upper bound can only be achieved by adding the newly scanned item to the items in the prior scan history. Formal definitions were given in ***Analysis of visual search behavior*** (set *T*_1_ and *T*_2_ respectively). We used |β_1_ | to measure the neuronal sensitivities to changes in the upper bound. A Signed-rank test was used to assess whether the slopes were significantly different across the recorded population. Neural activities were z-scored before linear regressions.

#### Upper bound neurons

We first computed the average impulse rate during each upper bound level by counting the number of impulses during each upper bound level and dividing the impulse counts by the time interval between start of the item scan the led to the current upper bound and the start of the item scan that led to the next upper bound level. To identify upper bound neurons, we then calculated the mean impulse rate for each upper bound level and correlated it with the upper bound using Spearman’s correlation (p < 0.05). To make sure that upper bound coding was not a simple consequence of movement preparations, we first regressed the neural impulse rate against the first movement directions and the time stamps when a new upper bound was achieved:

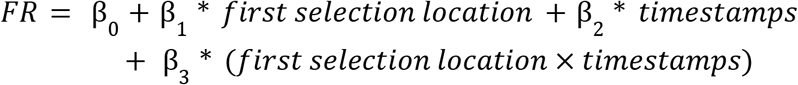

This formulation accounts for potentially different neural ramping corresponding to arm movements made to one of the six item locations. We then computed the Spearman’s correlation between the upper bound levels and their corresponding mean residuals.

To estimate the single-trial neural dynamics of single neurons, we obtained smoothed, single-trial impulse rate by applying kernel function (1 − *e*^−*t*^) * *e*^−*t*/30^ to the binned neural impulses (1ms bin). For each trial, we were interested in the pre-trial baseline-subtracted, estimated impulse rates between 100ms after the start of first item scan and first selections (pre-trial baseline was taken 1000ms before the onset of the instance). The evolution of the single-trial impulse rates as a linear function:

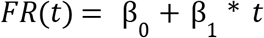

where β_1_ is the neural buildup rate. We then regressed the trial-by-trial neural buildup rate against the rate of upper bound growth, which was defined as the ratio between the final upper bound value during deliberation and the time interval between the start of the first item scan and the first selection. For hypothesis testing, we computed the trial-by-trial Spearman’s correlation between rate of upper bound growth and neural buildup rate within each neuron. The correlations were negated for neurons that showed negative upper bound coding, since faster upper bound growth should be associated with more negative β_1_ for those neurons. Finally, we used a Signed-rank test to assess whether the correlations were significantly different than 0.

#### Neural variability and behavioral performance

To assess whether neural variability is predictive of the animals’ behaviors on a trial-by-trial basis, we first derived the predicted impulse rate corresponding to the final upper bound value for each trial during deliberation with the following linear model:

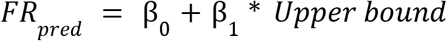

For each item scan during the final upper bound of each trial *t* (*u*_*t*_), we calculated the average impulse rate between 0 to 250ms since each item scan onset and selected the item scan with the maximum impulse rate for positive-coding upper bound neurons (and the minimum impulse rate for negative-coding upper bound neurons) as *FR*(*u*_*t*_). For each trial t, we then computed the neural deviation as

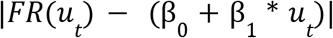

Finally, for each neuron, we calculated the Spearman’s correlation between the neural deviations and the rewards the animals obtained on individual trials.

### Statistics and Reproducibility

All statistical analyses were performed, and all graphs were created in MATLAB R2024b. No statistical methods were used to predetermine sample sizes, but our sample sizes are similar to those reported in studies investigating neuronal and behavioral responses in nonhuman primates. For inclusion of single units, we included well-isolated single units with more than 150 trials (202/208 neurons). All one-sample and two-sample statistical tests are nonparametric and two-sided. Effects were considered significant at *p* < 0.05. We used Benjamini-Hochberg Procedure to control False Discovery Rate when needed. Data collection and analysis were not performed blind to the conditions of the experiments.

---

**Fig. S1.**
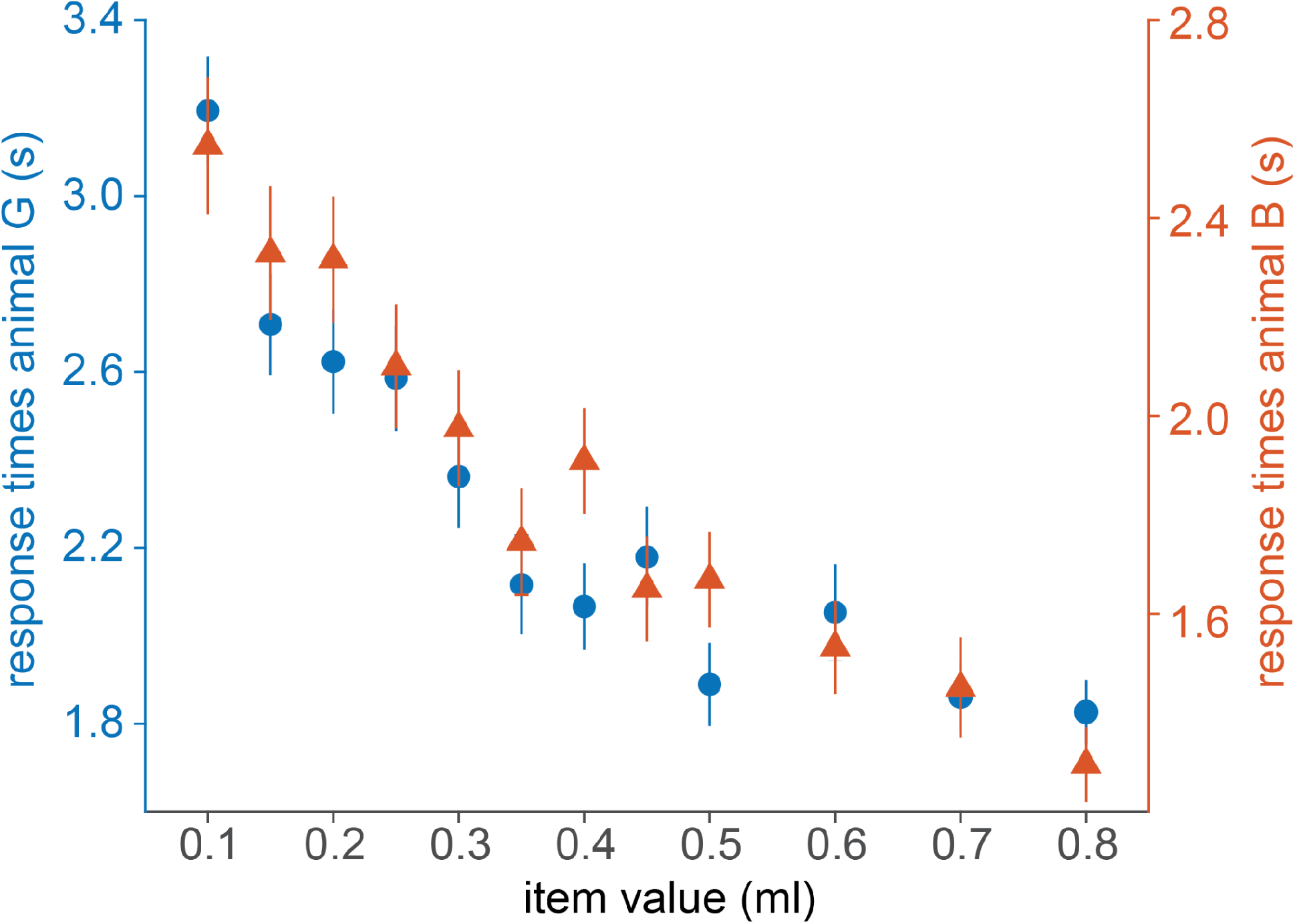
Animals learned the values of the individual cues. Scatter plot of response times vs reward magnitude for animal G (filled dots, left y-axis) and animal B (triangles, right y axis). Lower response times for large-reward cues indicated that the animals had learned the values. Error bars are SEM across trials of individual reward magnitudes.

**Fig. S2.**
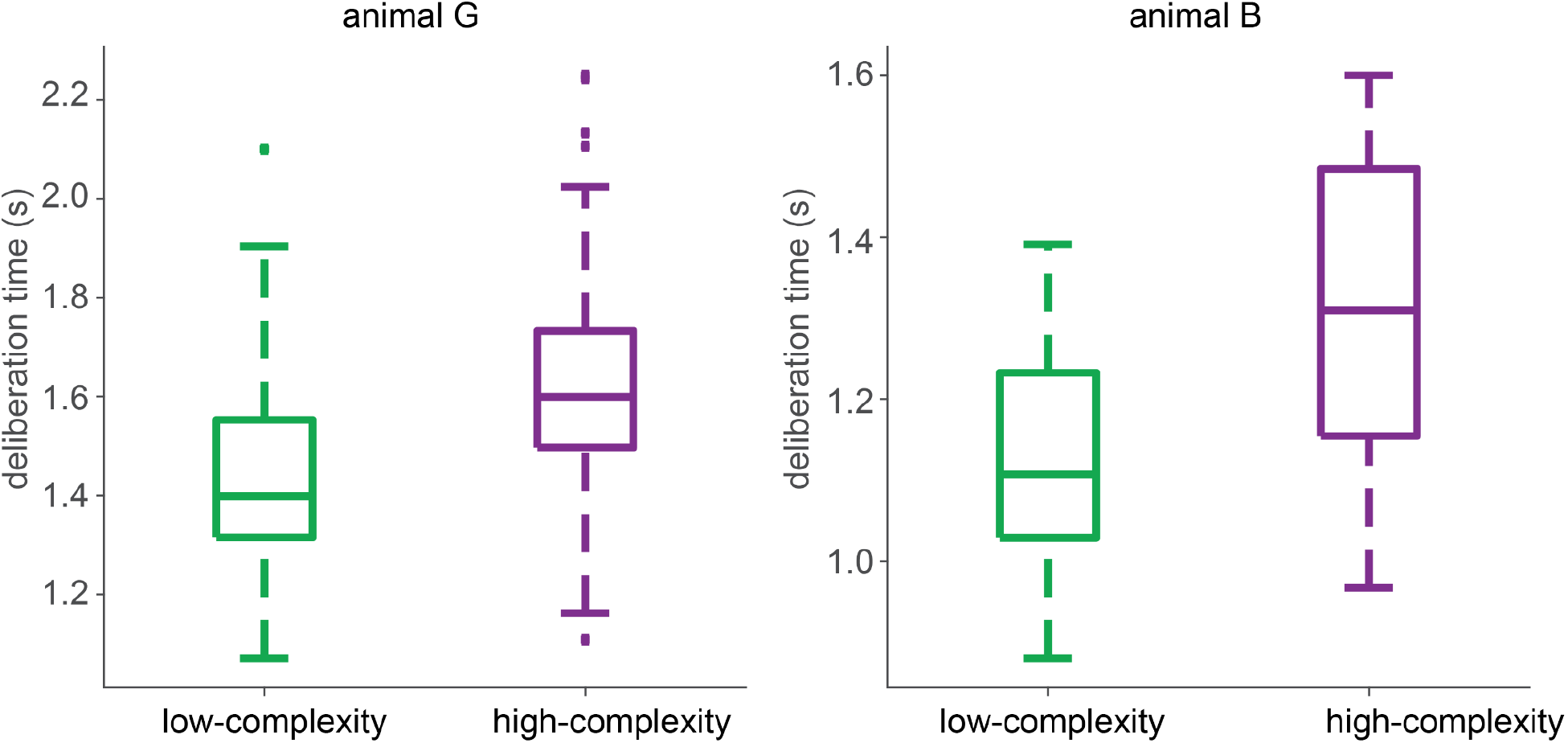
Deliberation times for low- and high-complexity trials. **(A)** Boxplots showing the deliberation times for low- (green) and high-complexity (purple) solution trials from animal G (p < 10^−14^, Signed-rank test). **(B)** Deliberation times for animal B (p < 10^−3^, Signed-rank test).

**Fig. S3.**
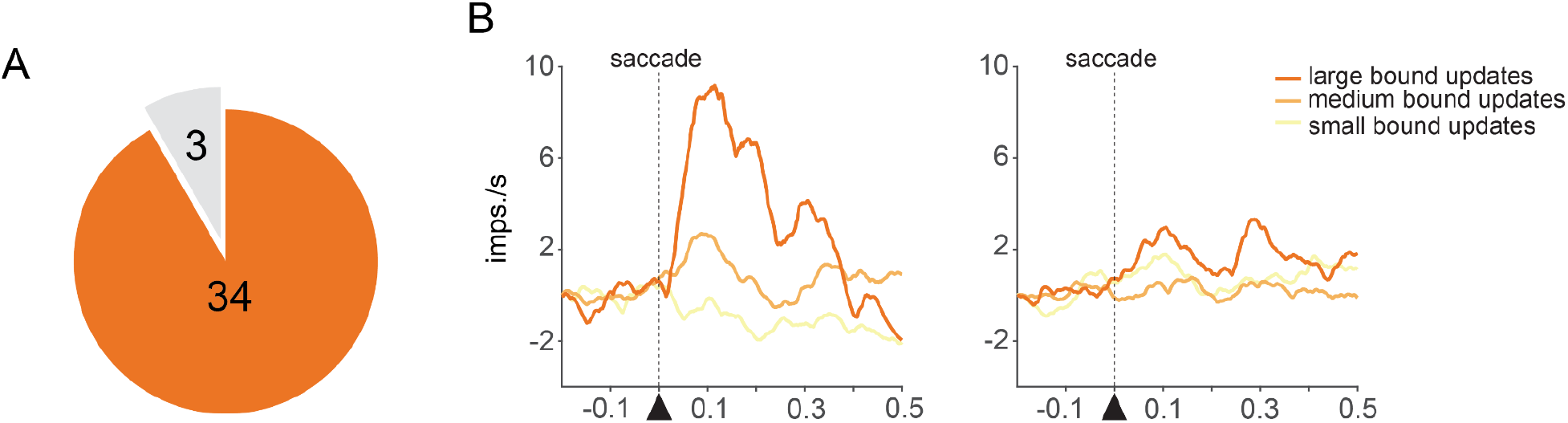
Only a small subset of bound-update neurons was modulated by saccade directions. **(A)** Pie chart showing the number of bound-update neurons that were modulated by saccade directions (grey) versus those that were not (orange, p > 0.05, two-way ANOVA). **(B)** Peri-event spike histogram (PETH) of an example bound-update neuron that was modulated by saccade directions. PETH on the left showed bound-update tuning when upper-right saccades led to bound updates. PETH on the right showed bound-update tuning for all other saccades that led to bound updates. Bound update magnitudes were binned into three bins (<=0.2ml, >0.2ml & <0.5ml, >=0.5ml).

## Notes

### Competing Interest Statement

The authors have declared no competing interest.

## References and Notes

1. S. Arora, B. Barak, Computational Complexity: A Modern Approach (Cambridge University Press, 2009).

2. P. Bossaerts, C. Murawski, Computational Complexity and Human Decision-Making. Trends in Cognitive Sciences 21, 917–929 (2017).

3. P. S. Goldman-Rakic, Cellular basis of working memory. Neuron 14, 477–485 (1995).

4. E. K. Miller, J. D. Cohen, An integrative theory of prefrontal cortex function. Annu. Rev. Neurosci. 24, 167–202 (2001).

5. S. M. Szczepanski, R. T. Knight, Insights into Human Behavior from Lesions to the Prefrontal Cortex. Neuron 83, 1002–1018 (2014).

6. J. M. Fuster, G. E. Alexander, Neuron activity related to short-term memory. Science 173, 652–654 (1971).

7. S. Funahashi, C. J. Bruce, P. S. Goldman-Rakic, Mnemonic coding of visual space in the monkey’s dorsolateral prefrontal cortex. J Neurophysiol 61, 331–349 (1989).

8. R. H. Bauer, J. M. Fuster, Delayed-matching and delayed-response deficit from cooling dorsolateral prefrontal cortex in monkeys. J. Comp. Physiol. Psychol. 90, 293–302 (1976).

9. D. J. Freedman, M. Riesenhuber, T. Poggio, E. K. Miller, Categorical representation of visual stimuli in the primate prefrontal cortex. Science 291, 312–316 (2001).

10. T. Yang, M. N. Shadlen, Probabilistic reasoning by neurons. Nature 447, 1075–1080 (2007).

11. A. Nieder, J. Freedman David, K. Miller Earl, Representation of the Quantity of Visual Items in the Primate Prefrontal Cortex. Science 297, 1708–1711 (2002).

12. P. Viswanathan, A. Nieder, Differential impact of behavioral relevance on quantity coding in primate frontal and parietal neurons. Curr. Biol. 25, 1259–1269 (2015).

13. A. Ramirez-Cardenas, M. Moskaleva, A. Nieder, Neuronal Representation of Numerosity Zero in the Primate Parieto-Frontal Number Network. Current Biology 26, 1285–1294 (2016).

14. X. Cai, C. Padoa-Schioppa, Contributions of orbitofrontal and lateral prefrontal cortices to economic choice and the good-to-action transformation. Neuron 81, 1140–1151 (2014).

15. T. Hosokawa, S. W. Kennerley, J. Sloan, J. D. Wallis, Single-neuron mechanisms underlying cost-benefit analysis in frontal cortex. J. Neurosci. 33, 17385–17397 (2013).

16. M. R. Roesch, C. R. Olson, Impact of expected reward on neuronal activity in prefrontal cortex, frontal and supplementary eye fields and premotor cortex. J Neurophysiol 90, 1766–1789 (2003).

17. R. E. Passingham, S. P. Wise, The Neurobiology of the Prefrontal Cortex: Anatomy, Evolution, and the Origin of Insight (OUP Oxford, 2012).

18. T. Hong, W. R. Stauffer, Computational complexity drives sustained deliberation. Nat. Neurosci. 26, 850–857 (2023).

19. A. H. Land, A. G. Doig, An automatic method of solving discrete programming problems. Econometrica 28, 497 (1960).

20. P. J. Kolesar, A branch and bound algorithm for the knapsack problem. Manage. Sci. 13, 723–735 (1967).

21. S. Martello, D. Pisinger, P. Toth, Dynamic programming and strong bounds for the 0-1 Knapsack Problem. Manage. Sci. 45, 414–424 (1999).

22. S. Martello, D. Pisinger, P. Toth, New trends in exact algorithms for the 0–1 knapsack problem. Eur. J. Oper. Res. 123, 325–332 (2000).

23. T. L. Patterson, S. Goldman, C. L. McKibbin, T. Hughs, D. V. Jeste, UCSD Performance-Based Skills Assessment: Development of a New Measure of Everyday Functioning for Severely Mentally Ill Adults. Schizophr. Bull. 27, 235–245 (2001).

24. C. R. Bowie, W. W. Leung, A. Reichenberg, M. M. McClure, T. L. Patterson, R. K. Heaton, P. D. Harvey, Predicting schizophrenia patients’ real-world behavior with specific neuropsychological and functional capacity measures. Biol. Psychiatry 63, 505–511 (2008).

25. J. Li, J. S. Skinner, K. McGarry, L. H. Nicholas, S.-P. Wang, E. Bollens-Lund, A. S. Kelley, Declines in Wealth Among US Older Adults at Risk of Dementia. JAMA Neurol. 80, 1250–1252 (2023).

26. C. Roan Gresenz, J. M. Mitchell, B. Rodriguez, R. S. Turner, H. W. van der Klaauw, The financial consequences of undiagnosed memory disorders (2024). 10.2139/ssrn.4852312.

27. S. Becker, M. Bode, K. Brockmann, T. Gasser, K. Michaelis, S. Solbrig, H.-C. Nuerk, C. Schulte, W. Maetzler, M. Zimmermann, D. Berg, I. Liepelt-Scarfone, Cognitive-driven activities of daily living impairment as a predictor for dementia in Parkinson disease: A longitudinal cohort study: A longitudinal cohort study. Neurology 99, e2548–e2560 (2022).

28. C. Padoa-Schioppa, J. A. Assad, Neurons in the orbitofrontal cortex encode economic value. Nature 441, 223–226 (2006).

29. B. Lau, P. W. Glimcher, Value representations in the primate striatum during matching behavior. Neuron 58, 451–463 (2008).

30. W. Schultz, P. Dayan, P. R. Montague, A neural substrate of prediction and reward. Science 275, 1593–1599 (1997).

31. N. Eshel, M. Bukwich, V. Rao, V. Hemmelder, J. Tian, N. Uchida, Arithmetic and local circuitry underlying dopamine prediction errors. Nature 525, 243–246 (2015).

32. P. Waelti, A. Dickinson, W. Schultz, Dopamine responses comply with basic assumptions of formal learning theory. Nature 412, 43–48 (2001).

33. E. E. Steinberg, R. Keiflin, J. R. Boivin, I. B. Witten, K. Deisseroth, P. H. Janak, A causal link between prediction errors, dopamine neurons and learning. Nat. Neurosci. 16, 966–973 (2013).

34. P. Bossaerts, W. Schultz, The neuroeconomics of simple and complex choice, Social Science Research Network (2025). 10.2139/ssrn.5352877.

35. K. Kubota, H. Niki, Prefrontal cortical unit activity and delayed alternation performance in monkeys. J Neurophysiol 34, 337–347 (1971).

36. G. B. Dantzig, Discrete-Variable Extremum Problems. Operations Research 5, 266–277 (1957).

37. E. Horowitz, S. Sahni, Computing partitions with applications to the knapsack problem. J. ACM 21, 277–292 (1974).

38. D. S. Johnson, Approximation algorithms for combinatorial problems. Journal of Computer and System Sciences 9, 256–278 (1974).

39. C. Murawski, P. Bossaerts, How Humans Solve Complex Problems: The Case of the Knapsack Problem. Scientific Reports 6, 34851 (2016).

40. C. J. Bruce, M. E. Goldberg, M. C. Bushnell, G. B. Stanton, Primate frontal eye fields. II. Physiological and anatomical correlates of electrically evoked eye movements. J Neurophysiol 54, 714–734 (1985).

